# The Druggable Genome as Seen from the Protein Data Bank

**DOI:** 10.1101/2020.02.06.937730

**Authors:** Jiayan Wang, Setayesh Yazdani, Ana Han, Matthieu Schapira

## Abstract

Almost twenty years after the human genome was sequenced, the wealth of data produced by the international human genome project has not translated into a significantly improved drug discovery enterprise. This is in part because small molecule modulators that could be used to explore the cellular function of their target proteins and to discover new therapeutic opportunities are only available for a limited portion of the human proteome. International efforts are underway to develop such chemical tools for a few, specific protein families, and a “Target 2035” call to enable, expand and federate these efforts towards a comprehensive chemical coverage of the druggable genome was recently announced. But what is the druggable genome? Here, we systematically review structures of human proteins bound to drug-like ligands available from the protein databank (PDB) and use ligand desolvation upon binding as a druggability metric to draw a landscape of the human druggable genome. We show that the vast majority of druggable protein families, including some highly populated and deeply associated with cancer according to genomic screens, are almost orphan of small molecule ligands, and propose a list of 46 druggable domains representing 3440 human proteins that could be the focus of large chemical probe discovery efforts.

## Introduction

Continuous advances in library screening technologies such as DNA-encoded libraries (Neri and Lerner, 2018), fragment screening (Erlanson et al., 2016) or free energy perturbation (Wang et al., 2015a) enable the discovery of drug-like small molecule ligands for an increasing number of protein targets. In parallel, the boundary of target druggability is constantly evolving, sometimes via a precise understanding of the structural dynamics of a ligand binding site (Kessler et al., 2019), or via the emergence of novel therapeutic modalities (Lai and Crews, 2017). These progresses and future technological advances are expected to enable a significantly expanded chemical coverage of the human proteome.

Born from both this favorable context and also the realization that chemical probes can play a decisive role in the discovery of novel therapeutic targets (Arrowsmith et al., 2015; Oprea et al., 2018), large-scale efforts are ongoing to increase the chemical coverage of specific protein families, such as protein kinases, solute carriers, GPCRs and epigenetic target classes (César-Razquin et al., 2015; Knapp et al., 2012; Roth et al., 2017; Scheer et al., 2019; Wu et al., 2019). Based on the progress with these efforts, a vision is emerging to create chemical ligands for the entire druggable genome by year 2035 (Carter et al., 2019). Delineating what is the druggable genome will help guide this ambitious enterprise (Finan et al., 2017; Hopkins and Groom, 2002; Nguyen et al., 2017; Oprea et al., 2018; Russ and Lampel, 2005; Santos et al., 2017).

Here, we systematically analyze protein domains that are in complex with drug-like ligands in the Protein Data Bank (PDB) to evaluate the druggability and structural coverage of target classes in the human genome. While restricted to protein families with bound ligands in the PDB, this approach, firmly anchored in experimental data, largely recapitulates previous classifications, but also reveals emerging target classes, some highly populated, that could be the focus of future chemical biology and drug discovery efforts.

### The chemical coverage of the human proteome is poor

We first extracted all drug-like ligands from the PDB and identified the Interpro domain of the protein to which each ligand was bound. For each domain, we next counted the number of different proteins in which the domain was bound to a drug-like ligand in the PDB (Supplementary Tables 1 and 2). Unsurprisingly, we found that kinases are the most ligand-bound protein family (195 proteins bound to 2091 ligands in 2536 structures). Far behind were GPCRs (37 proteins bound to 67 ligands in 98 structures), followed by nuclear hormone receptors (25 proteins bound to 515 ligands in 686 structures) and bromodomains (22 proteins bound to 297 ligands in 365 structures). The chemical coverage (as seen from the PDB) of bromodomains is however greater than that of kinases: 54% of bromodomains, 52% of nuclear hormone receptors and 40% of kinases are bound to a drug-like ligand (Fig. 1). This number drops to 5% for GPCRs, but this reflects the difficulty to crystallize these proteins and their limited presence in the PDB rather than the extent of their chemical coverage. Similarly, ion channels structures typically are solved at resolutions above 3.0 Å, which is the cut-off for this analysis. As a result, this important class of drug targets (Hopkins and Groom, 2002; Oprea et al., 2018; Santos et al., 2017) is under-represented in our survey. Together, these data indicate that vast areas of target classes known to be druggable, such as kinases or nuclear hormone receptors remain underexplored. The fact that over 75% of protein families composed of 30 or more proteins have less than 10% of their members in complex with drug-like ligands in the PDB clearly illustrates the paucity of the chemical coverage of the human genome (Fig. 1, Supplementary Table 1).

**Figure 1:**
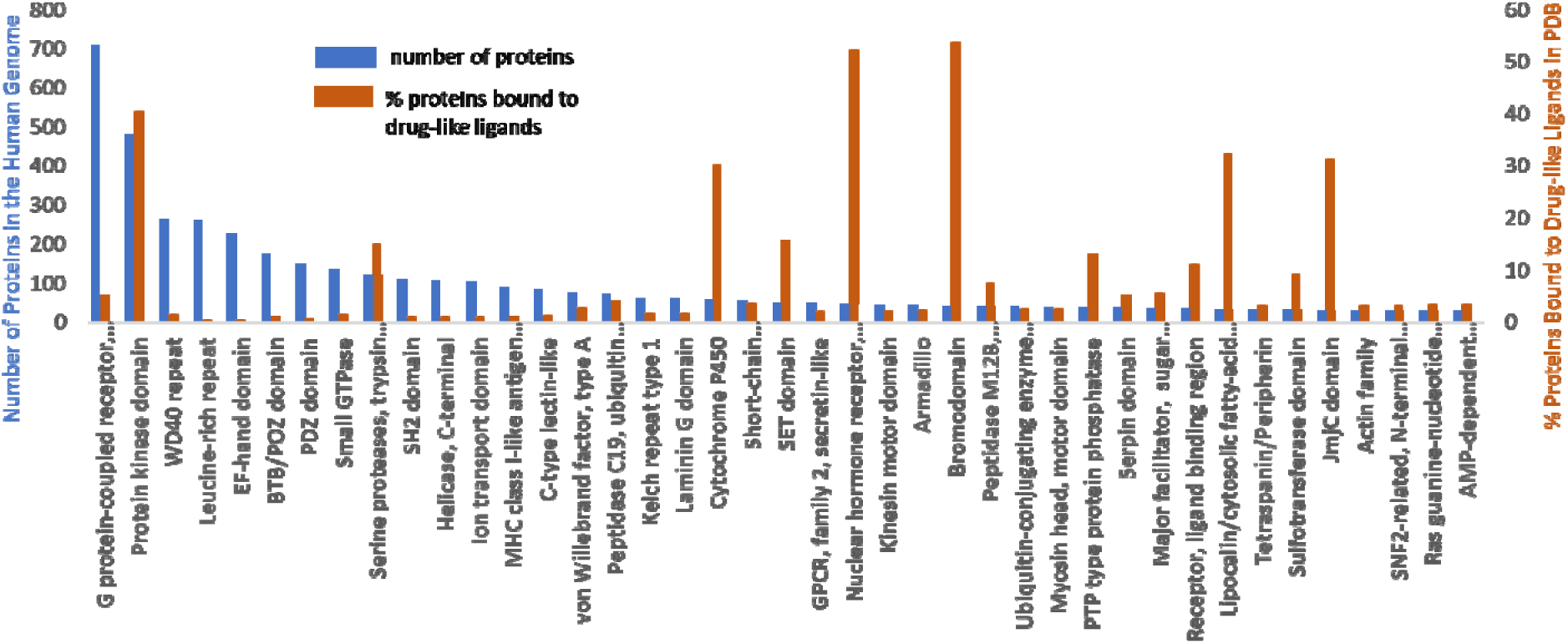
Chemical coverage in the PDB of the most frequent protein domains found in the human genome. Only protein domains with at least one structure in the PDB bound to a drug-like ligand with a resolution of 3Å or better are included in the analysis. Proteins domains found in less than 30 human proteins are not shown.

### Some large and druggable protein families in the Protein Data Bank are almost entirely devoid of drug-like ligands

We note that some of the largest protein families have very few ligands bound. The WDR family is predicted to contain up to 610 proteins (Wang et al., 2015b), but only 4 WDR domains are bound to drug-like ligands in the PDB. Only 1 leucine-rich repeat (LRR) and 1 EF-hand containing protein are bound to a ligand, but each family includes over 200 genes. To evaluate whether lack of druggability is underlying this deficiency in chemical coverage, we estimated the druggability of protein families with two orthogonal approaches. In the first method, we calculated the percent surface of the ligand that is desolvated when bound to the protein. This value reflects the buriedness of the ligand in the pocket, and is expected to correlate with the druggability of the protein. The second method relies on the PDBbind database, which compiles the affinity of some of the ligands found in the PDB for their bound proteins (Wang et al., 2005). When multiple ligands were available for a protein, we used the strongest affinity and highest desolvation state, and the median value across all proteins was used for each domain. We purposely omitted the use of geometrical or electrostatic features of pockets to predict druggability, as is often done (Borrel et al., 2015; Halgren, 2009), including by us (Song et al., 2017), because we set out to avoid any computational prediction in this analysis. We saw a correlation between the two metrics: protein families with high median ligand desolvation generally have low median ligand affinity values (i.e. measured binding is stronger). Small GTPases are an exception, as structures of KRAS bound to potent inhibitors were recently deposited in the PDB, leading to high desolvation values, but these ligands are not yet entered in the PDBbind database, where only weaker KRAS ligands are recorded (Supplementary Table 2).

We find that nuclear hormone receptors, cytochrome P450s and GPCRs are among the most druggable target classes, all having median ligand desolvation values of 95% or more, and affinity values (when available) below 100 nM (Fig. 2). LRR seem to position at the other end of the druggability spectrum, with a desolvation value of 57% (Supplementary Table 2). Though largely overlooked by the drug discovery and chemical biology community, the WDR family - one of the most populated in the human genome – seem eminently druggable, with a high median ligand desolvation of 77% and a low median affinity of 48 nM (Fig. 2, Supplementary Table 2). These data are derived though from only 4 targets (EED, WDR5, CDC20 and BTRC), and extrapolation to the entire protein family is uncertain, but it is nevertheless an encouraging indicator. WDR are β-propeller domains with a canonical toroidal shape (Schapira et al., 2017), and reported ligands exploit the central cavity of EED (He et al., 2017; Qi et al., 2017), WDR5 (Grebien et al., 2015) and BTRC (Simonetta et al., 2019) with low nanomolar potency, and the side wall of CDC20 (Sackton et al., 2014) with micromolar affinity.

**Figure 2:**
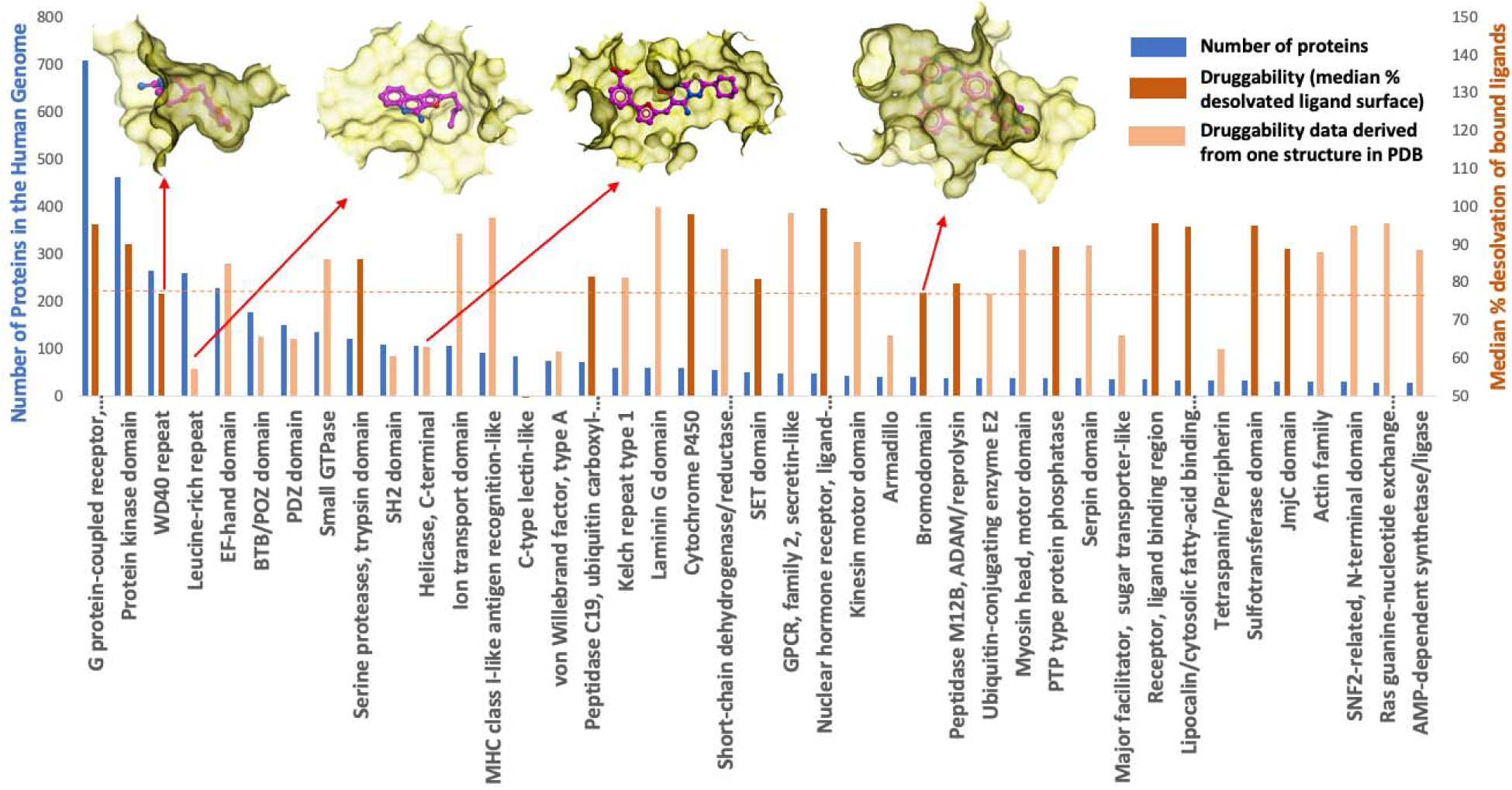
Druggability of target classes. For each protein family, the median desolvation value of bound ligands across all proteins in the PDB is used as an indicator of druggability (the most desolvated ligand is used for each protein). Dashed line: median desolvation of bromodomains. Faded orange color indicates families where the druggability data is based on less than three proteins (mean values are used instead of median). Crystal structures of ligand-bound pockets are shown for four representative domains: two with high ligand desolvation (WDR [PDB: 5u5t], Bromodomain [PDB: 4o76]) and two with low ligand desolvation (Leucine-rich repeat [PDB: 3wn4], helicase [PDB: 5urm]).

### WDR proteins are a large, druggable and underexplored target class with strong disease association

To evaluate the disease association of severely underexplored but putatively druggable target classes, such as WDRs, and compare it with that of protein families more popular in drug discovery, we analyzed the essentiality of proteins across multiple cancer types based on CRISPR-knockout data available from the Broad and the Wellcome Sanger Institutes (Behan et al., 2019; Meyers et al., 2017). For each cancer type represented by at least 3 cell lines, the median essentiality score across all cell lines was determined for each gene, and the number of genes essential (essentiality score <-1.) in at least 3 cancer types was determined for each protein family (Fig. 3). Strikingly, we found that the WDR family is populated with the highest number of essential genes (Broad: 54 genes; Sanger: 64 genes), followed by helicases (Broad: 22 genes; Sanger: 32 genes) and protein kinases (Broad: 14 genes; Sanger: 20 genes). While helicases are low on our druggability ladder (the only ligand in the PDB is 63% desolvated when bound to its target, SNRNP200 [PDB code 5urm]), available data indicates that WDRs are druggable (see above), and the exceptional disease association of this target classes is a call for increased efforts to develop small-molecule WDR antagonists.

**Figure 3:**
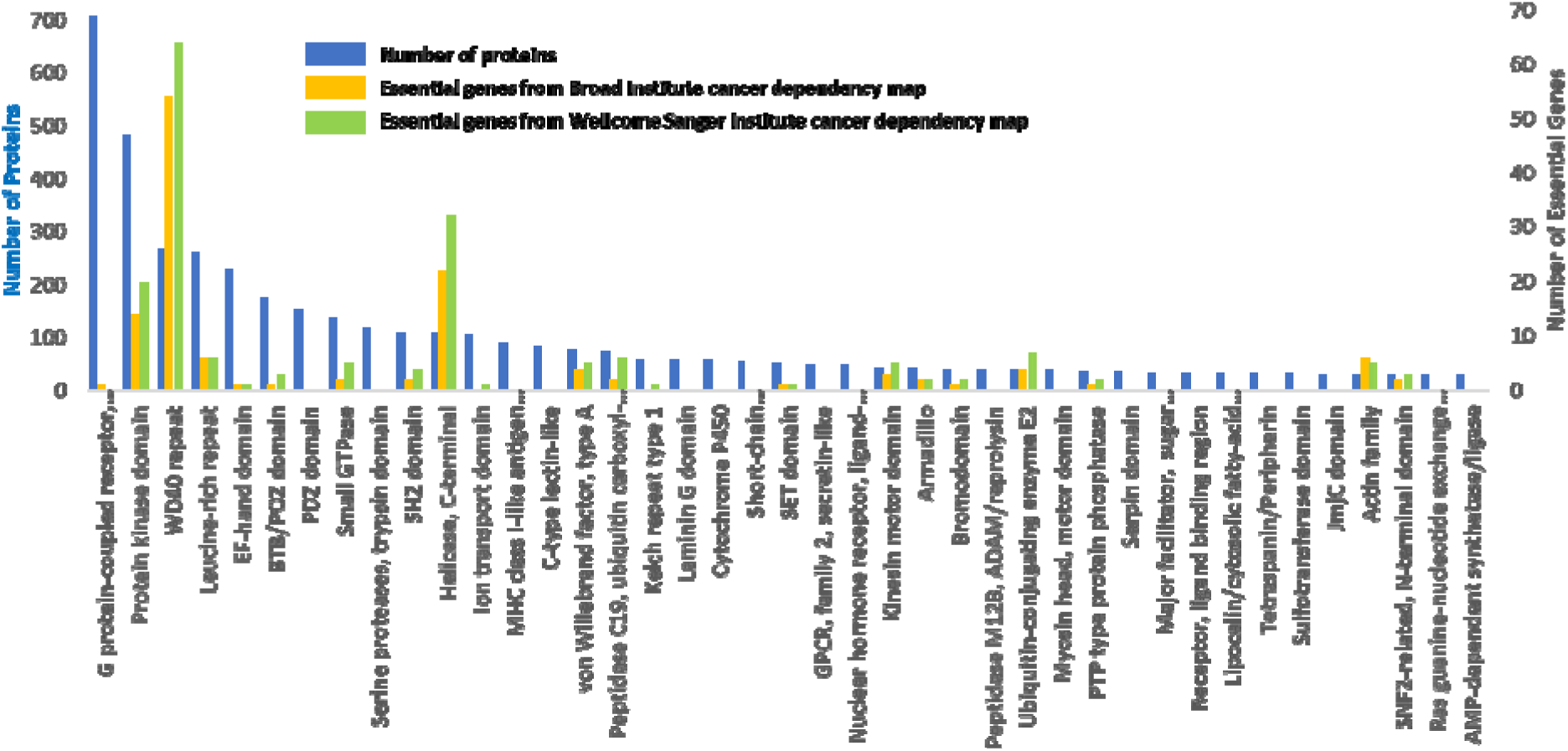
Essentiality of protein families in cancer. The number of proteins essential in at least 3 cancer types based on cancer dependency maps from the Broad and the Wellcome Sanger Institutes are indicated (details provided in the supplementary information).

### Structural data currently in the PDB suggests that a large number of protein families are more druggable than bromodomains in the human genome

Overall, we find that 163 protein families have a median ligand desolvation level greater than that of bromodomains (desolvation value: 77%), a druggable target class (Supplementary Table 2). Of these, 46 are populated with 20 or more proteins, and could be the object of a focused medicinal chemistry effort where knowledge acquired and ligands discovered for one protein may guide and accelerate the design of ligands for another, as was done for kinases (Elkins et al., 2016) or bromodomains (Wu et al., 2019). We believe that these 46 protein families (Table 2) representing 3440 proteins (Supplementary Table 3) could be a good place to start a global effort towards the discovery of chemical ligands for the druggable proteome (Carter et al., 2019).

**Table 2:**
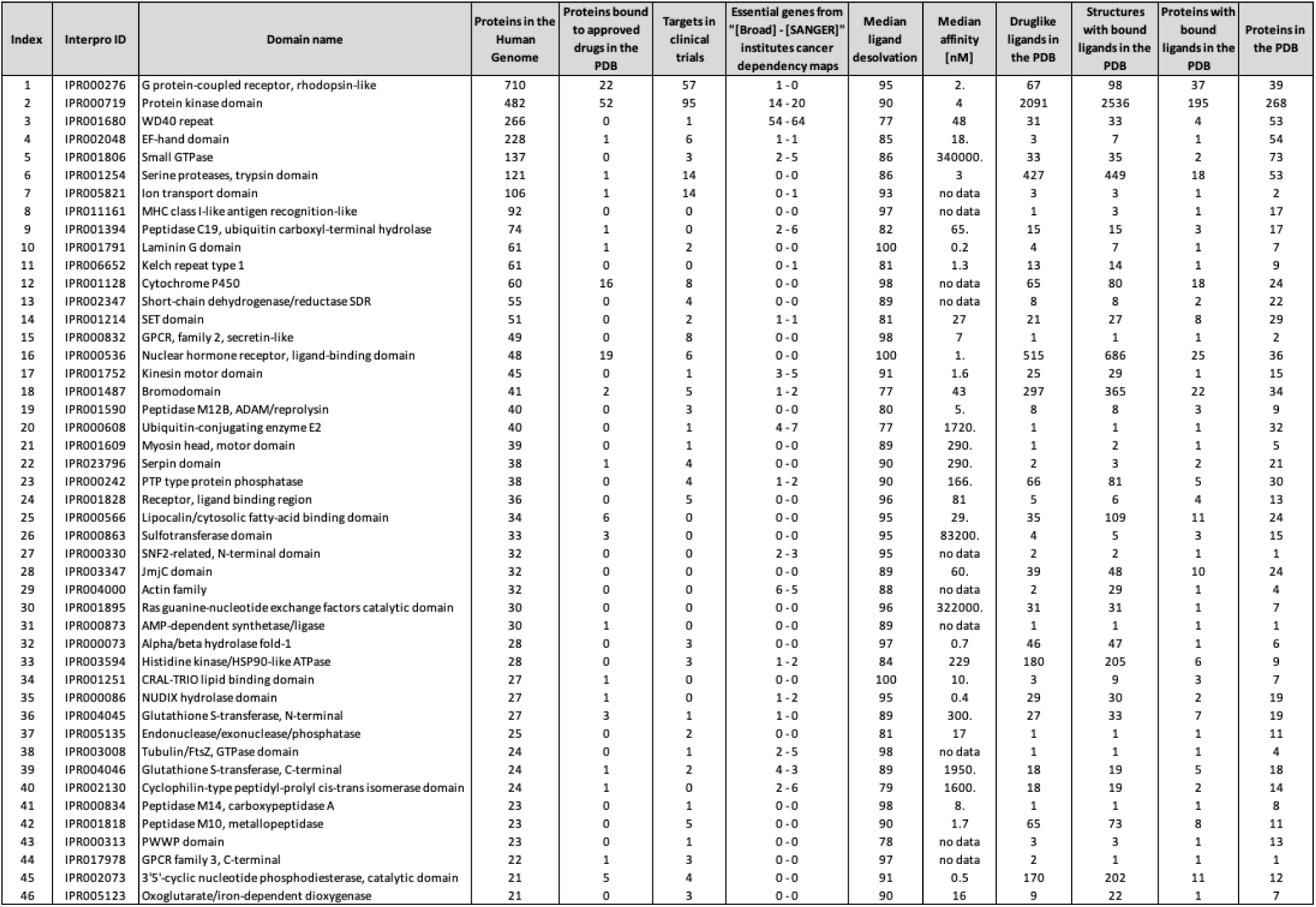
Human druggable genome, represented by protein families of 20 or more proteins, with at least one protein bound to a drug-like inhibitor in the PDB (with a resolution better than 3 Å), and median ligand desolvation upon binding equal or better than that of bromodomains.

## Conclusion

Motivated by the principle that experimental structures of drug-like ligands in complex with their receptor proteins are among the most reliable data linking ligands and protein domains, we provide here an analysis of the druggable genome as seen from the PDB. We find that a significant portion of well-known druggable target classes remain underexplored, and that other druggable protein families, some large and with strong disease association, are almost entirely orphan of inhibitors. The list of 3440 proteins from 46 druggable target classes compiled here may help strategize emerging efforts to systematically develop chemical reagents for the human proteome (Carter et al., 2019).

## Acknowledgments

The SGC is a registered charity (number 1097737) that receives funds from AbbVie, Bayer Pharma AG, Boehringer Ingelheim, Canada Foundation for Innovation, Eshelman Institute for Innovation, Genome Canada through Ontario Genomics Institute [OGI-055], Innovative Medicines Initiative (EU/EFPIA) [ULTRA-DD grant no. 115766], Janssen, Merck KGaA, Darmstadt, Germany, MSD, Novartis Pharma AG, Innovation and Science (MRIS), Pfizer, São Paulo Research Foundation-FAPESP, Takeda, and Wellcome. M.S. gratefully acknowledges support from NSERC [Grant RGPIN-2019-04416]. We thank Aled Edwards for comments on the manuscript.

