## Supplementary information for "The Druggable Genome as Seen from the Protein Data Bank"

**Supplementary data**

**Supplementary table 1:** Chemical coverage of protein domains as seen from the PDB. Details in caption to Fig. 1

**Supplementary table 2**: dynamic table available at <http://polymorph.sgc.utoronto.ca/drugged_human_proteome/>

with links to all data associated to all protein domains presented in this work, including interactive 3D images of all ligands bound to all proteins. The web interface is also permanently archived at <https://doi.org/10.5281/zenodo.3645838>

**Supplementary table 3**: list of 46 protein families and their 3440 human proteins that are at least as druggable as bromodomains.

**Supplementary Methods**

All scripts and data were generated with Molsoft’s ICM and are permanently available at <http://doi.org/10.5281/zenodo.3653135>. These files can be opened with Molsoft’s free ICM Browser (www.molsoft.com).

**Extracting drug-like ligands from the PDB:**

30280 compounds were downloaded from the Protein Data Bank. Compounds with negative drug-likeness (calculated with ICM [Molsoft, San Diego]) were discarded. The Lipinski rule of five was applied (no more than one violation of the following four criteria: MW<500 Da, cLogP<5, H-bond donors <=5, H-bond acceptors <= 10). Additional hard limits were set as follows: 200 Da<MW<700 Da, cLogP>-3, H-bond donors <=10, H-bond acceptors <= 20. Ligands with phosphate groups and ligands without a ring or with more than five fused rings were also discarded. Lastly, 11 sugar structures were removed. The finak set contained 11419 drug-like ligands.

**Mapping ligands to protein domains:**

16346 human protein structures bound to drug-like ligands with resolution better than 3 Å were found in the PDB. For each structure, the biologically relevant oligomeric state was generated with ICM. Residues within 4 Å of the ligand were used to define the sequence boundaries of the binding site. Binding pockets formed by less than 7 residues, or located at the interface of multiple protein chains, were discarded. The binding pocket sequence was then mapped onto the reference human proteome from Uniprot with Blast using 95% sequence identity cut-off. The resulting sequence encompassing the ligand-binding pocket was then cross-referenced against protein domain boundaries available from the InterPro database where 90% of the query or the hit should be covered. The domain was considered valid if at least 4 amino acid residues were involved in ligand interactions. Because of a high level of redundancy in InterPro, we limited our domain definition to InterPro domains sourced from the Pfam database. The Serine-Threonine/Tyrosine kinase domain (Interpro IPR001245) was merged with the protein kinase domain (Interpro IPR000719). 7742 ligand-bound pockets were mapped to 258 proteins domains.

**Ligand desolvation**:

For each ligand-bound pocket, the percentage of the ligand surface area accessible to solvent was calculated with ICM (command: “show surface area”, followed by “Acc(a_lig//*)”. See supplementary ICM scripts).

**Gene essentiality in cancer:**

We used data from the cancer dependency maps at Sanger (Behan et al, Nature 2019) and Broad (Meyers et al. Nat Genet. 2017) Institutes that were derived from CRISPR-cas9 knockout experiments in various cancer cell lines. We considered a gene to be essential in a given cancer type if the cancer type was represented by at least 3 cell lines and if the median essentiality score was lower <-1.

**Approved Drugs**:

ICM (Molsoft, San Diego) was used to identify all druglike ligands from our list with chemical structures identical to approved drugs from DrugBank (Wishart et al Nucleic Acids Res. 2018) (Tanimoto fingerprint distance < 0.02; non-exact matches were deleted manually). By cross-referencing the list of Uniprot domains, and their corresponding proteins and bound ligands in the PDB with the list of matched approved drugs, approved drugs were mapped to different protein families. In the end, the total number of proteins targeted by approved drugs for each protein family was reported.

**Clinical Trials**:

The list of current targets in clinical trials from the Therapeutic Target Database (Wang et al. Nucleic Acids Res. 2020) were matched to the list of gene names in their corresponding protein families. In the end, the total number of targets in clinical trials for each protein family was reported.
