## Supplementary Table 1 for "The Druggable Genome as Seen from the Protein Data Bank"

|  | Inerpro ID | Domain Name | Proteins in the Human Genome | Drug-like ligands in PDB | Structures with bound ligand in PDB | Proteins with Bound Ligand in PDB |
| --- | --- | --- | --- | --- | --- | --- |
| 1 | IPR000276 | G protein-coupled receptor, rhodopsin-like | 710 | 67 | 98 | 37 |
| 2 | IPR000719 | Protein kinase domain | 482 | 2091 | 2536 | 195 |
| 3 | IPR001680 | WD40 repeat | 266 | 31 | 33 | 4 |
| 4 | IPR001611 | Leucine-rich repeat | 261 | 2 | 2 | 1 |
| 5 | IPR002048 | EF-hand domain | 228 | 3 | 7 | 1 |
| 6 | IPR000210 | BTB/POZ domain | 178 | 7 | 7 | 2 |
| 7 | IPR001478 | PDZ domain | 152 | 1 | 1 | 1 |
| 8 | IPR001806 | Small GTPase | 137 | 33 | 35 | 2 |
| 9 | IPR001254 | Serine proteases, trypsin domain | 121 | 427 | 449 | 18 |
| 10 | IPR000980 | SH2 domain | 109 | 3 | 3 | 1 |
| 11 | IPR001650 | Helicase, C-terminal | 108 | 1 | 1 | 1 |
| 12 | IPR005821 | Ion transport domain | 106 | 3 | 3 | 1 |
| 13 | IPR011161 | MHC class I-like antigen recognition-like | 92 | 1 | 3 | 1 |
| 14 | IPR001304 | C-type lectin-like | 86 | 1 | 1 | 1 |
| 15 | IPR002035 | von Willebrand factor, type A | 76 | 7 | 7 | 2 |
| 16 | IPR001394 | Peptidase C19, ubiquitin carboxyl-terminal hydrolase | 74 | 15 | 15 | 3 |
| 17 | IPR006652 | Kelch repeat type 1 | 61 | 13 | 14 | 1 |
| 18 | IPR001791 | Laminin G domain | 61 | 4 | 7 | 1 |
| 19 | IPR001128 | Cytochrome P450 | 60 | 65 | 80 | 18 |
| 20 | IPR002347 | Short-chain dehydrogenase/reductase SDR | 55 | 8 | 8 | 2 |
| 21 | IPR001214 | SET domain | 51 | 21 | 27 | 8 |
| 22 | IPR000832 | GPCR, family 2, secretin-like | 49 | 1 | 1 | 1 |
| 23 | IPR000536 | Nuclear hormone receptor, ligand-binding domain | 48 | 515 | 686 | 25 |
| 24 | IPR001752 | Kinesin motor domain | 45 | 25 | 29 | 1 |
| 25 | IPR000225 | Armadillo | 42 | 3 | 3 | 1 |
| 26 | IPR001487 | Bromodomain | 41 | 297 | 365 | 22 |
| 27 | IPR001590 | Peptidase M12B, ADAM/reprolysin | 40 | 8 | 8 | 3 |
| 28 | IPR000608 | Ubiquitin-conjugating enzyme E2 | 40 | 1 | 1 | 1 |
| 29 | IPR001609 | Myosin head, motor domain | 39 | 1 | 2 | 1 |
| 30 | IPR000242 | PTP type protein phosphatase | 38 | 66 | 81 | 5 |
| 31 | IPR023796 | Serpin domain | 38 | 2 | 3 | 2 |
| 32 | IPR005828 | Major facilitator, sugar transporter-like | 36 | 2 | 2 | 2 |
| 33 | IPR001828 | Receptor, ligand binding region | 36 | 5 | 6 | 4 |
| 34 | IPR000566 | Lipocalin/cytosolic fatty-acid binding domain | 34 | 35 | 109 | 11 |
| 35 | IPR018499 | Tetraspanin/Peripherin | 33 | 1 | 1 | 1 |
| 36 | IPR000863 | Sulfotransferase domain | 33 | 4 | 5 | 3 |
| 37 | IPR003347 | JmjC domain | 32 | 39 | 48 | 10 |
| 38 | IPR004000 | Actin family | 32 | 2 | 29 | 1 |
| 39 | IPR000330 | SNF2-related, N-terminal domain | 32 | 2 | 2 | 1 |
| 40 | IPR001895 | Ras guanine-nucleotide exchange factors catalytic domain | 30 | 31 | 31 | 1 |

**Supplementary Table 1: Chemical coverage of protein domains as seen from the PDB**.
